# Partial or complete loss of norepinephrine differentially alters contextual fear and catecholamine release dynamics in hippocampal CA1

**DOI:** 10.1101/2023.03.26.534277

**Authors:** Leslie R. Wilson, Nicholas W. Plummer, Irina Y. Evsyukova, Daniela Patino, Casey L. Stewart, Kathleen G. Smith, Sydney A Fry, Alex L. Deal, Victor W. Kilonzo, Natale R. Sciolino, Jesse D. Cushman, Patricia Jensen

**Affiliations:** Neurobiology Laboratory, National Institute of Environmental Health Sciences, National Institutes of Health, Dept. of Health and Human Services, Research Triangle Park, NC, USA; Neurobehavioral Core Laboratory, National Institute of Environmental Health Sciences, National Institutes of Health, Dept. of Health and Human Services, Research Triangle Park, NC, USA; Dept. of Physiology and Neurobiology, Dept. of Biomedical Engineering, Institute for System Genomics, Connecticut Institute for the Brain & Cognitive Sciences, University of Connecticut, Storrs, CT, USA

## Abstract

Contextual fear learning is heavily dependent on the hippocampus. Despite evidence that catecholamines contribute to contextual encoding and memory retrieval, the precise temporal dynamics of their release in the hippocampus during behavior is unknown. In addition, new animal models are required to probe the effects of altered catecholamine synthesis on release dynamics and contextual learning. Utilizing GRAB_NE_ and GRAB_DA_ sensors, *in vivo* fiber photometry, and two new mouse models of altered locus coeruleus norepinephrine (LC-NE) synthesis, we investigate norepinephrine (NE) and dopamine (DA) release dynamics in dorsal hippocampal CA1 during contextual fear conditioning. We report that aversive foot-shock increases both NE and DA release in dorsal CA1, while freezing behavior associated with recall of fear memory is accompanied by decreased release. Partial loss of LC-NE synthesis reveals that NE release dynamics are modulated by sex. Moreover, we find that recall of recent fear memory is sensitive to both partial and complete loss of LC-NE synthesis throughout prenatal and postnatal development, similar to prior observations of mice with global loss of NE synthesis beginning postnatally. In contrast, remote recall is compromised only by complete loss of LC-NE synthesis beginning prenatally. Overall, these findings provide novel insights into the role of NE in contextual fear and the precise temporal dynamics of both NE and DA during freezing behavior, and highlight a complex relationship between genotype, sex, and NE signaling.

## INTRODUCTION

Contextual learning involves the integration of the multisensory features of a particular environment into a single representation that can support complex adaptive behaviors (1–3). Individual stimuli can have substantially different meaning and significance in different contexts, and contexts themselves can serve as powerful predictors of appetitive and aversive events. In contextual fear conditioning, the multisensory features that define the conditioning chamber are associated with an aversive unconditional stimulus that subsequently drives fear expression (3–6). Contextual fear is heavily dependent on the hippocampus (7, 8) and numerous studies have demonstrated the importance of norepinephrine (NE) and dopamine (DA) signaling in different hippocampal subregions to contextual encoding and retrieval (9–16).

With respect to NE, the hippocampus receives virtually all of its inputs from the locus coeruleus (LC; (17, 18)), and LC-NE signaling to dorsal hippocampal area CA1 (dCA1) has been specifically implicated in retrieval of contextual memories (19, 20). LC-NE signaling through β-adrenergic receptors has been shown to modulate activation of dCA1 pyramidal neurons during memory retrieval (11), and optogenetic activation of LC-NE inputs to dCA1 enhances retrieval of contextual memories 24 hours after training (21). Consistent with these observations, global loss of NE synthesis in dopamine β-hydroxylase knockout mice (*Dbh^ko^*; (22)) results in deficits in short-term memory retrieval (11). Despite these important discoveries, the precise temporal dynamics of NE and DA release in dCA1 during fear conditioning remain unknown. In addition, it is unclear how catecholamine dynamics and contextual learning are altered upon more subtle disruptions of NE synthesis.

In the current study we present two new mouse models of disrupted LC-NE synthesis beginning prenatally: a hypomorphic allele of *Dbh* resulting in reduced NE in central noradrenergic neurons and an LC conditional knockout with total loss of NE in the LC and reduced NE in other central noradrenergic centers. Utilizing these mice in conjunction with a pre-exposure-dependent contextual fear assay (23, 24) and *in vivo* fiber photometry in dCA1, we determine the effects of partial and complete embryonic loss of LC-NE synthesis on contextual memory, and assess NE and DA release dynamics during four phases of contextual learning. Our data identify deficits in memory retrieval and unique patterns of catecholamine release dynamics in dCA1 during memory acquisition and retrieval that are differentially sensitive to partial or complete embryonic loss of NE.

## RESULTS

### Generation of a mouse model of disrupted LC NE synthesis

In previous studies using *Dbh^ko^* mice, the embryonic lethality believed to be caused by loss of NE synthesis in the peripheral nervous system was overcome by dosing pregnant dams with L-*threo*-3,4-dihydroxyphenylserine (L-DOPS), permitting *Dbh*-independent NE synthesis during prenatal development (22, 25). Consequently, *Dbh^ko^* mice lack global NE postnatally, after L-DOPS treatment is withdrawn. While the *Dbh^ko^* is an excellent model to assess the requirement for NE throughout the postnatal period, the restoration of NE synthesis mediated by L-DOPS may obscure phenotypes associated with prenatal loss of NE. To eliminate LC-NE synthesis beginning in embryonic development and avoid the prenatal lethality associated with global loss of *Dbh*, we generated a Cre-dependent conditional knockout allele, *Dbh^tm2.2Pjen^* (*Dbh^flx^*), in which fluorescent tags allow the recombination status of individual noradrenergic neurons to be monitored. We inserted a loxP site within intron 1 and replaced the stop codon in exon 12 with a construct consisting of a viral 2A peptide (26), tdTomato expression cassette, loxP site, splice acceptor sequence, and EGFP expression cassette (Fig. 1A). In the absence of Cre-mediated recombination, transcription from the *Dbh^flx^* allele will produce an mRNA encoding full-length DBH and tdTomato. Due to the presence of the 2A peptide sequence, translation will result in separate DBH and tdTomato proteins. Cre-mediated recombination will delete most of the *Dbh* coding sequence, leaving a mutant allele (*Dbh^null^*) that encodes a fusion between exon 1 and EGFP. Thus, if Cre is expressed in a subset of noradrenergic neurons, the Cre+ noradrenergic neurons can be identified by EGFP fluorescence, while Cre-negative noradrenergic neurons that retain DBH activity are labeled with tdTomato. To confirm that recombination occurs as expected, we crossed *Dbh^flx^* heterozygotes with *Tmem163^Tg(ACTB-cre)2Mrt^* mice (27) which exhibit ubiquitous Cre activity. Polymerase chain reaction (PCR) analysis of genomic DNA demonstrated Cre-dependent deletion of sequence between the two loxP sites (Fig. 1A).

**Figure 1.**
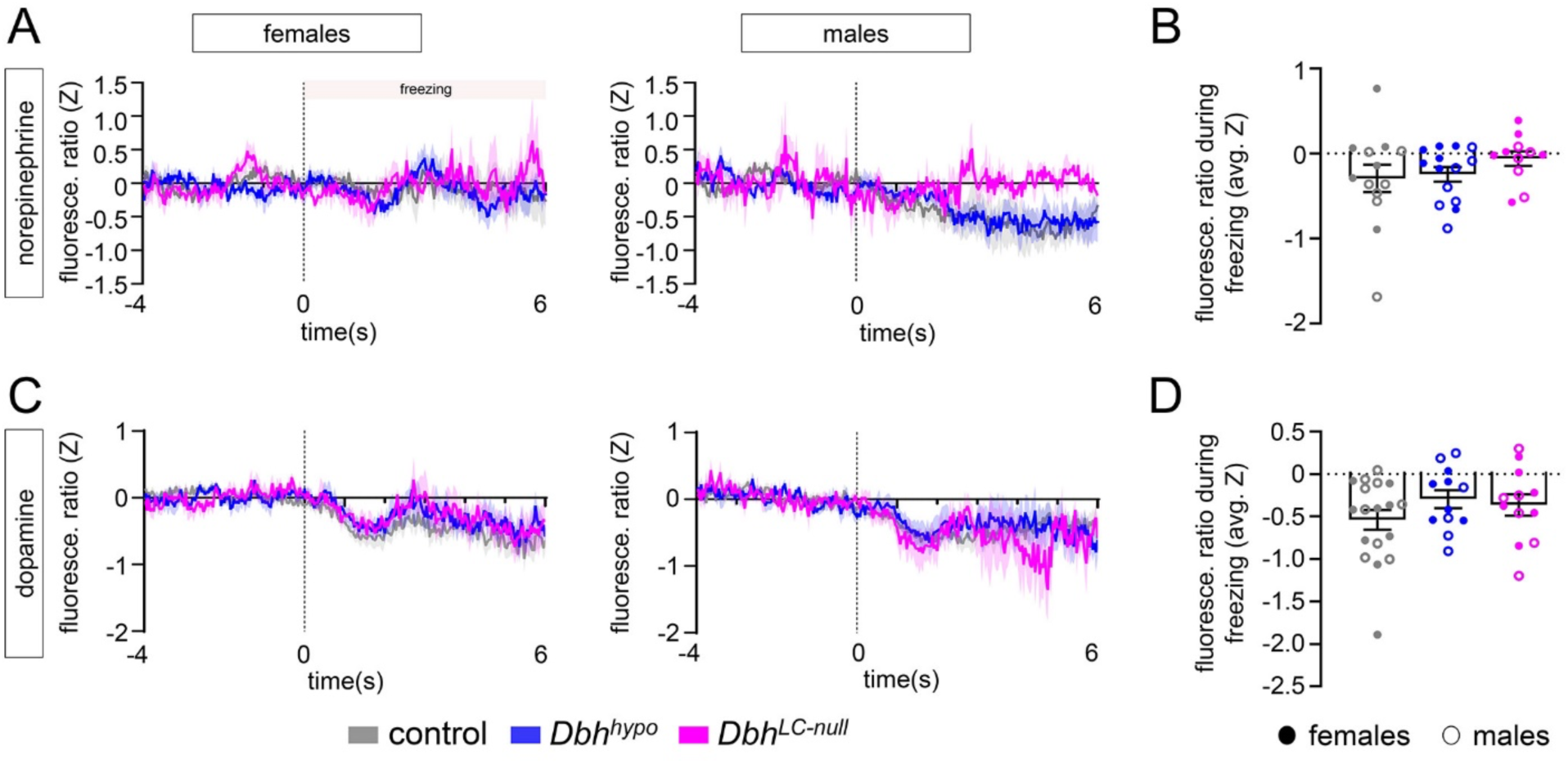
Conditional knockout allele of the mouse dopamine β-hydroxylase (*Dbh*) gene. (**A**) Schematic diagram showing the modified *Dbh* locus. Coding exons are indicated by black boxes, 5’ and 3’ untranslated regions by white boxes, and introns by black lines. In the conditional (*Dbh^flx^*) allele, a single loxP site is inserted within intron 1, and a larger insertion in exon 12 consists of 2A peptide (26), tdTomato cDNA, SV40 polyadenylation signal, loxP, splice acceptor sequence (s.a.), EGFP cDNA, and rabbit β-globin poly adenylation signal. After Cre recombination, the null allele consists of exon 1, a chimeric intron consisting of the 5’ portion of intron 1 and the exogenous splice acceptor, and EGFP. Arrows indicate PCR primers flanking the loxP sites. Box: PCR analysis of genomic DNA from *Dbh^wt/wt^* (wild type), heterozygous *Dbh^flx/wt^*, homozygous *Dbh^flx/flx^*, and heterozygous *Dbh^null/wt^* mice demonstrates excision of exons 2-12 and the tdTomato cassette following Cre recombination. (**B**) Sagittal schematic diagram of the embryonic neural tube showing overlap of noradrenergic progenitors and *En1* expression domain in the anterior hindbrain, and adult sagittal schematic showing location of noradrenergic nuclei and expected fluorescent labeling of noradrenergic neurons in *Dbh^LC-null^* (*En1^Cre/wt^ Dbh^flx/flx^*). (**C**) Representative sections from a Cre-negative *Dbh^flx/flx^* and *Dbh^LC-null^* mouse showing tdTomato or EGFP expression in tyrosine hydroxylase (TH)-expressing noradrenergic neurons of the locus coeruleus (LC), dorsal subcoeruleus (SubCD), ventral subcoeruleus (SubCV), C2/A2, and C1/A1 nuclei. Scale bar: 100 µm.

Next, we selectively eliminated NE synthesis in LC neurons by taking advantage of the fact that the anatomically defined LC (within the central gray of the pons, adjacent to the fourth ventricle), together with a rostral portion of the adjacent dorsal subcoeruleus, is uniquely defined by embryonic expression of the transcription factor *Engrailed 1* (*En1*) and later expression of *Dbh* (18). Because *En1* is expressed before *Dbh*, the LC neurons of mice heterozygous for *En1^Cre^* (28) and homozygous for *Dbh^flx^* (*Dbh^LC-null^* mice; Fig. 1B) will never produce DBH protein, and therefore never synthesize NE. As expected, LC noradrenergic neurons of *Dbh^LC-null^* mice were labeled with EGFP, indicating loss of *Dbh* expression (Fig. 1C). In non-LC noradrenergic neurons, we observed tdTomato, indicating the unrecombined *Dbh* allele (Fig. 1C). Detection of both EGFP and tdTomato required immunofluorescent labeling; nevertheless, these results indicate that the fluorescent tags can be used to monitor recombination status of the *Dbh^flx^* allele.

### Mutation in *Dbh* intron 1 enhancer is associated with partial disruption of NE synthesis

To confirm that *Dbh^LC-null^* mice lack *Dbh* expression in the LC, we performed co-immunofluorescent labeling of DBH and tyrosine hydroxylase (TH). Consistent with the recombination data (Fig. 1C), DBH was not detected in the LC of *Dbh^LC-null^* mice (Fig. 2A). We observed no differences in DBH labeling between Cre+ and Cre-negative *Dbh^wt/wt^* controls, and no genotype-specific differences in TH labeling. However, in Cre-negative *Dbh^flx/flx^* mice, DBH labeling appeared qualitatively weaker than in *Dbh^wt/wt^* controls (Fig. 2A).

**Figure 2.**
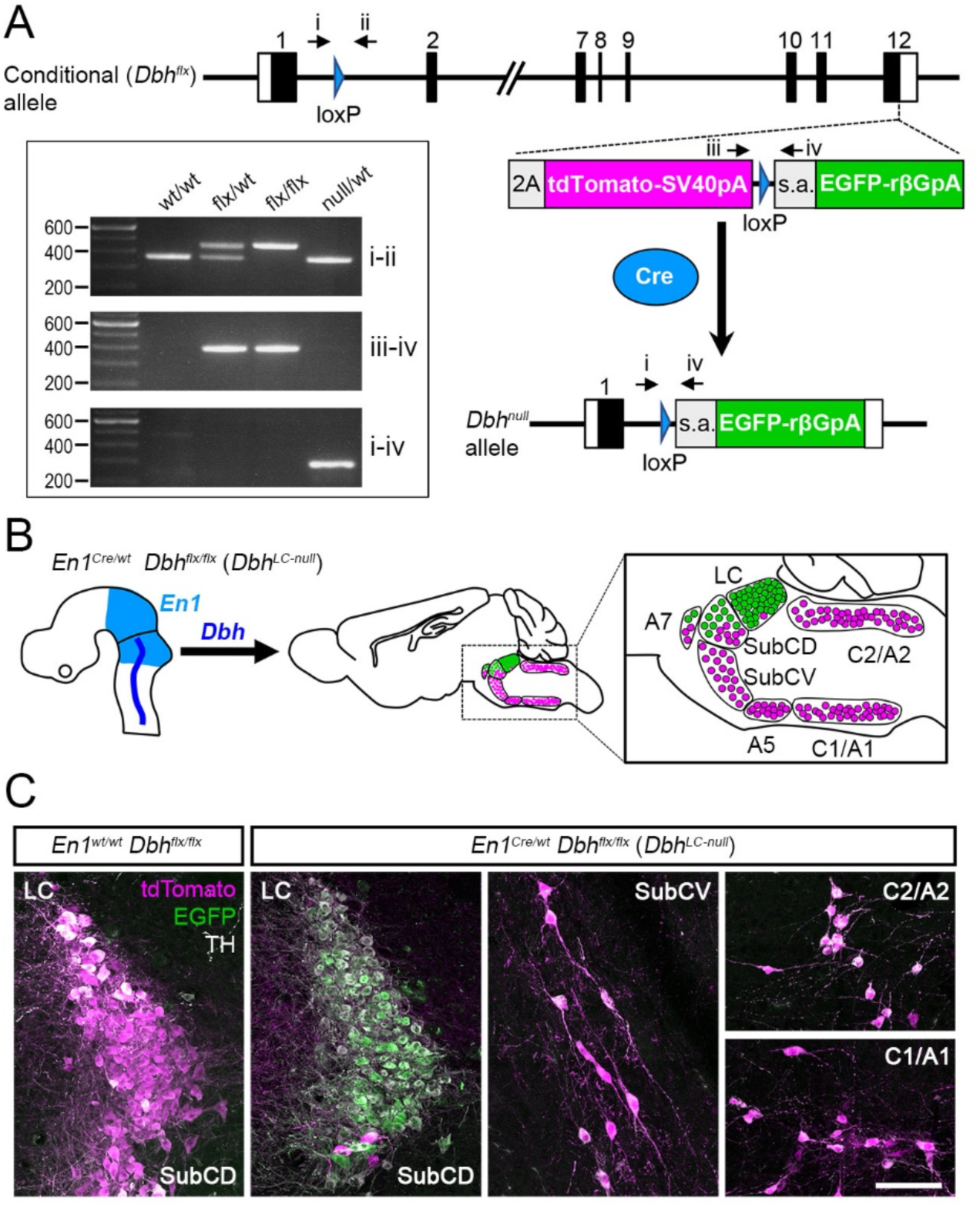
Disruption of NE synthesis in *Dbh^LC-null^* and *Dbh^flx/flx^* (*Dbh^hypo^*) mice. (**A**) Coronal sections from *Dbh^wt/wt^*, *Dbh^flx/flx^*, and *Dbh^LC-null^* mice showing DBH (green) and/or TH (magenta) expression in the LC (representative images from n=4 female and 4 male per genotype). Scale bar = 100 µm. (**B**) Levels of NE and DA or DOPAC in hippocampus and plasma of *Dbh^wt/wt^*, *Dbh^flx/flx^*, and *Dbh^LC-null^* mice. Data are mean ± SEM. For hippocampus samples, n = 7 F (closed circles) and 8 M (open circles) *Dbh^wt/wt^* mice (includes 8 *En1^Cre/wt^* and 7 *En1^wt/wt^*), 5 F and 5 M *Dbh^flx/flx^* mice, and 11 F and 13 M *Dbh^LC-null^* mice. For plasma samples, n = 14 F and 18 M *Dbh^wt/wt^* (17 *En1^Cre/wt^* and 15 *En1^wt/wt^*), 10 F and 9 M *Dbh^flx/flx^*, and 10 F and 10 M *Dbh^LC-null^*. ****p<0.0001. ^###^p=0.0004 (main effect of sex). **(C)** Levels of Dbh and Th mRNA in tissue samples containing noradrenergic neurons from *Dbh^wt/wt^* and *Dbh^flx/flx^* mice. Data are mean ± SEM. For *Dbh^flx/flx^* pons, n = 4 females (closed circles) and 3 males (open circles). For all other samples, n = 4 females and 4 males. ****p<0.0001, **p=0.0020.

To confirm that disruption of LC *Dbh* expression eliminates NE synthesis, we examined NE tissue content in the hippocampus using mass spectrometry. As expected, NE was absent in the hippocampus of *Dbh^LC-null^* mice (Fig. 2B). Consistent with DBH immunolabeling, we observed an approximately 60% reduction of hippocampal NE in Cre-negative *Dbh^flx/flx^* mice relative to *Dbh^wt/wt^ controls*, demonstrating that the *Dbh^flx^* allele is hypomorphic (Fig. 2B). We also found that NE content was significantly higher in *Dbh^wt/wt^* and *Dbh^flx/flx^* male mice compared to females (Fig. 2B; p=0.003, sex by genotype interaction). Because DBH converts DA to NE, we also examined DA content in the hippocampus and found that it was significantly increased in *Dbh^flx/flx^* and *Dbh^LC-null^* mice relative to *Dbh^wt/wt^* controls (Fig. 2B; p<0.0001, main effect of genotype, 2-way ANOVA with Tukey’s test). Furthermore, DA in *Dbh^LC-null^* hippocampus was significantly higher than in *Dbh^flx/flx^* (Fig. 2B; p<0.0001, 2-way ANOVA with Tukey’s test), demonstrating that larger disruptions in NE synthesis correspond to greater hippocampal DA levels. In contrast, we observed no differences between genotypes in plasma levels of the DA metabolite DOPAC or NE, which represents release from sympathetic neurons (29) indicating that NE release from the peripheral nervous system is unaffected in *Dbh^LC-null^* and *Dbh^flx/flx^* mice. Taken together with the analysis of recombination (Fig. 1), our findings confirm that *Dbh^LC-null^* mice lack NE synthesis in the LC, and additionally that the *Dbh^flx^* allele is hypomorphic rather than functionally wild type.

To explore possible causes of the unexpected hypomorphy of *Dbh^flx/flx^* mice (hereafter designated *Dbh^hypo^*), we examined regulatory annotations for the mouse genome (GRCm39) in the Ensembl Genome Browser (Release 108) (30). We found that a 1399 bp putative enhancer region in mouse *Dbh* intron 1 (Ensembl ENSMUSR00000824281) is interrupted by the upstream loxP insertion at an EcoRI site beginning at nucleotide 291. Disruption of an enhancer is expected to alter mRNA levels; therefore, we performed droplet digital PCR (ddPCR) to assess Dbh mRNA in central noradrenergic centers (Fig. 2C). We examined mRNA levels in pons (which includes LC, subcoeruleus and A5 nuclei) and medulla (A1 and A2 nuclei). In *Dbh^hypo^* mice, we found reduced Dbh mRNA relative to wild type in both regions (reduction of approximately 73% in pons, 64% in medulla). We also tested whether reduced *Dbh* expression and concomitant increase in DA levels (Fig. 2B) affects *Th* mRNA expression. We found no difference in *Th* mRNA expression between *Dbh^hypo^* and *Dbh^wt/wt^* samples (Fig. 2C), consistent with the TH immunolabeling in the LC (Fig. 2A). These data indicate that hypomorphic NE levels in *Dbh^hypo^* mice result from reduced *Dbh* mRNA synthesis. *Dbh^hypo^* and *Dbh^LC-null^* mice therefore represent two distinct models of disrupted NE synthesis: *Dbh^hypo^* with reduced NE in central noradrenergic neurons, and *Dbh^LC-null^* with total loss of NE in the LC and reduced NE in other noradrenergic centers.

### Impaired contextual fear response in *Dbh^hypo^* and *Dbh^LC-null^* mice

It has previously been shown that global loss of NE synthesis beginning in the early postnatal period results in deficits in hippocampal-dependent contextual fear memory (11). To assess the effects of embryonic loss of LC-NE—both partial and complete—on contextual memory and NE and DA release dynamics, we used spectrally resolved fiber photometry (31) combined with contextual fear conditioning. Male and female *Dbh^LC-null^*, *Dbh^hypo^* and littermate control mice were injected with AAVs expressing tdTomato and either a NE sensor (GRAB_NE_; (32)) or DA sensor (GRAB_DA_; (33)) and implanted with fiber optical probes in hippocampal dCA1(Fig. 3A). Next, we utilized the pre-exposure-dependent contextual fear assay (Fig. 3B), allowing four phases of contextual learning to be isolated. A pre-exposure phase allows the mice to explore the novel context and acquire a hippocampus-dependent contextual representation of the conditioning chamber. During the immediate-shock day this contextual representation is rapidly retrieved and associated with the shock to form a contextual fear memory. Recent (24 hours after shock) and remote (2 weeks after shock) tests assess short- and long-term retrieval of the contextual fear memory, as measured by the amount of time the mice spend freezing (i.e., motionless, a species-specific fear response) when placed back in the conditioning chamber.

**Figure 3.**
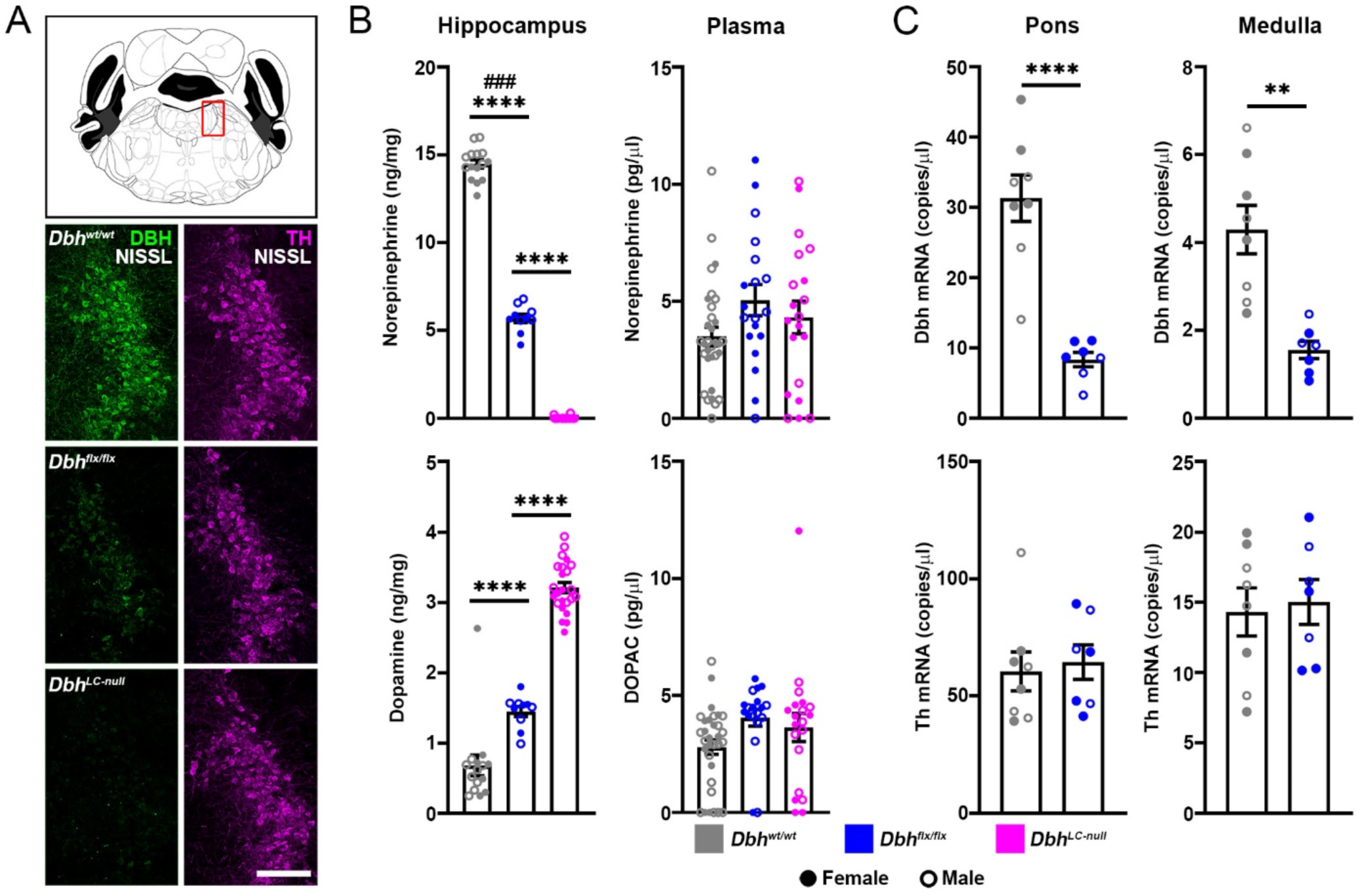
Experimental conditions and results of context-dependent fear conditioning. (**A**) Coronal schematic of mouse brain showing injection site of AAV9-GRAB_NE_ or -GRAB_DA_ and AAV5-tdTomato viruses in the dCA1 region of the dorsal hippocampus. Fiber optic cables were implanted three weeks later above the dCA1. *in vivo* fiber photometry recordings were performed using a spectrometer-based system with a 488nm emission laser. Software allowed for simultaneous visualization of photon counts from fluorescent signals at various wavelengths. (**B**) Schematic of context-dependent fear conditioning paradigm performed in parallel with fiber photometry. (**C**) Distance moved of the mice during the pre-exposure recording. (**D**) Distance moved during foot shock. (**E**) Percentage of time spent freezing during the recent context test (**, p=0021; *,p=0.031). (**F**) Freezing during the remote context test (**, p=0.0042). Data are mean ± SEM. 2-way ANOVA with Tukey’s multiple comparisons test, n=31 controls (15 F, 16 M), n=26 *Dbh^hypo^* (13 F, 13 M), and n=23 *Dbh^LC-null^* (12 F, 11 M).

During the pre-exposure phase of the test, we observed no genotype- or sex-specific differences in the distance moved, indicating no baseline differences motor exploration were present (Fig. 3C). Similarly, no genotype-specific differences in distance moved were observed during the activity burst associated with the two-second aversive foot shock (Fig. 3D). However, males were observed to move more than females (p=0.0090, main effect of sex). These results indicate that NE deficits associated with the *Dbh^hypo^* and *Dbh^LC-null^* mouse models do not affect overall activity or sensitivity to aversive stimuli. To assess retrieval of contextual fear memory we measured the percentage of time the mice spent freezing when placed back in the chamber in which they received the shock. During the short-term retrieval test, both *Dbh^hypo^* and *Dbh^LC-null^* mice exhibited reduced freezing compared to controls (Fig. 3E; *Dbh^hypo^*, p= 0.0305; *Dbh^LC-nul^*, p= 0.0021). However, during the remote retrieval test, only *Dbh^LC-null^* mice still exhibited significantly reduced freezing compared to controls (Fig. 3F; p= 0.0042). Taken together, these results indicate that recent contextual fear memory retrieval is sensitive to both partial and complete embryonic loss of LC NE, while remote contextual memory is sensitive only to complete loss.

### Dynamic release of NE and DA in dCA1 during contextual fear conditioning

Next, we analyzed the fiber photometry data to assess NE and DA dynamics in hippocampal dCA1 during the four phases of the fear conditioning paradigm. The photometry signal was aligned to the aversive shock and to the onset of freezing during the pre-exposure phase and tests of memory retrieval. During the pre-exposure, freezing was relatively rare but sufficient to permit analysis of event-triggered averages. We observed no genotype- or sex-specific differences in NE or DA release during freezing in the pre-exposure phase (Fig. S1), consistent with our behavioral observation that *Dbh^LC-null^*, *Dbh^hypo^* and littermate control mice have similar locomotion at baseline. However, during the aversive foot shock, we observed an immediate increase in NE release in both *Dbh^hypo^* and control mice. As expected, the evoked increase in NE was absent in *Dbh^LC-null^* mice (Fig. 4A-B), consistent with our measure of tissue content (Fig. 2B). During the 10 seconds following the foot shock, NE release was significantly increased in both wild type controls (p=0.0009) and *Dbh^hypo^* (p=0.0384) compared to *Dbh^LC-null^* mice (Fig. 4C). We also observed increased DA release in response to the shock (Fig. 4A,D), with release in *Dbh^LC-null^* mice significantly greater than in *Dbh^hypo^* (p=0.0067) and wild type controls (p<0.0001; Fig. 4E). Despite the significant difference in NE and DA tissue content between *Dbh^hypo^* and wild type mice (Fig. 2B), we found no significant differences in NE or DA release between those two genotypes (Fig. 4C, E). However, we did note a trend for a genotype by sex interaction for NE release (p=0.059 2-way ANOVA), in which male *Dbh^hypo^* mice resembled wild type males and female *Dbh^hypo^* resembled female *Dbh^LC-null^* (Fig 4B).

**Figure 4.**
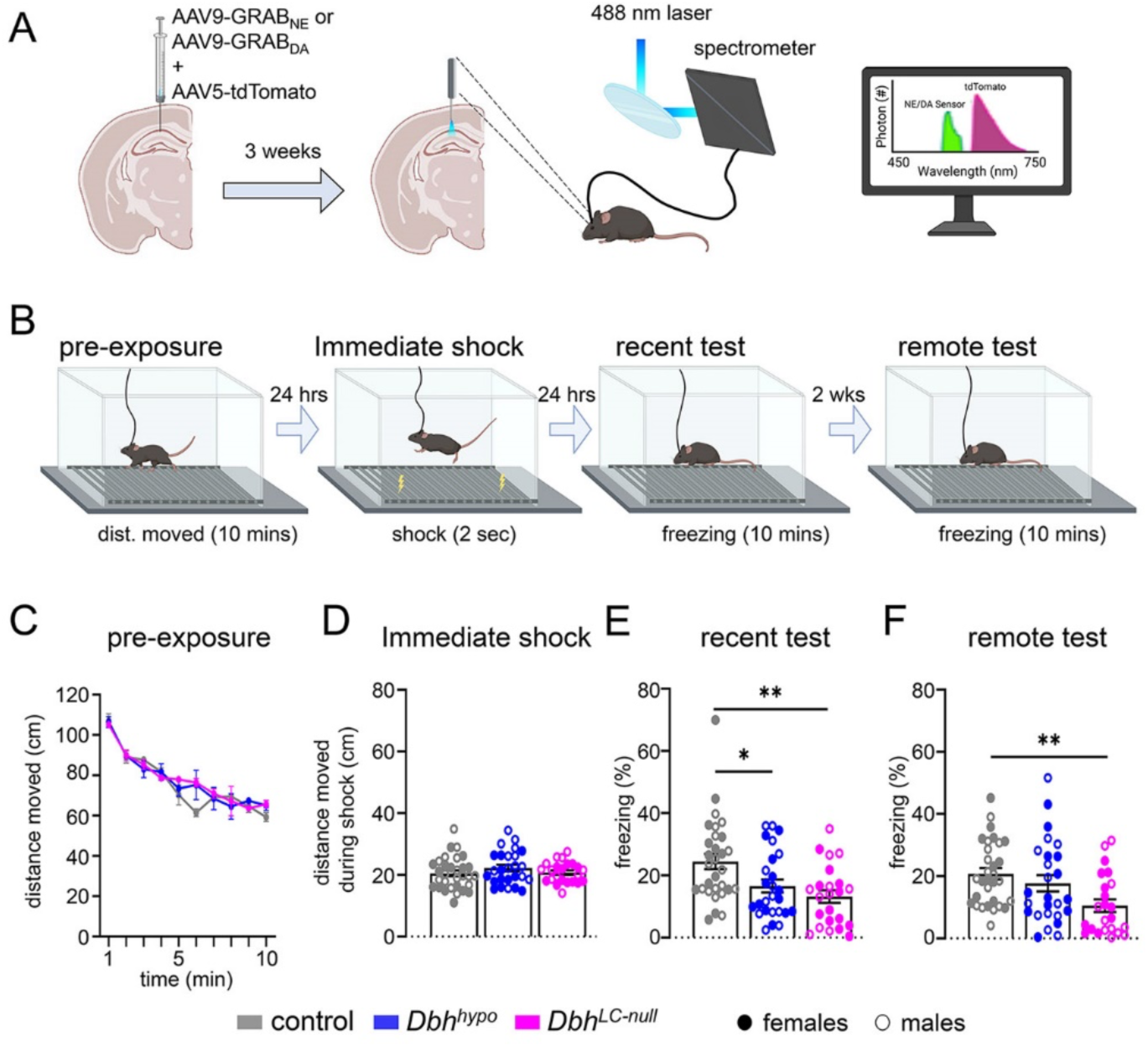
NE and DA dynamics in dCA1 during and immediately following a noxious foot shock. (**A**) GRAB_NE_/tdTomato (norepinephrine) and GRAB_DA_/tdTomato (dopamine) fluorescence ratios expressed as a normalized response aligned to foot shock (dotted line). Each row in the heat map corresponds to an individual mouse, with largest ratio for that mouse set to 100 and smallest ratio set to 0. (**B**) Average GRAB_NE_/tdTomato fluorescence ratios aligned to shock in female (left) and male (right) control, *Dbh^hypo^*, and *Dbh^LC-null^* mice. (**C**) Average NE response during the first ten seconds after the foot shock for the controls (95% CI of mean: lower, 1.31; upper, 4.76), *Dbh^hypo^* (lower, 1.03; upper, 2.83), and *Dbh^LC-null^* mice (lower, -0.64; upper, 0.56; encompasses zero). ***,p=0.0009; *, p=0.0384. (**D**) Average GRAB_DA_/tdTomato fluorescence ratios aligned to shock. (**E**) Average DA response during the first ten seconds after shock for the controls (95% CI of mean: lower, 0.98; upper, 2.30), *Dbh^hypo^* (lower, 1.10; upper, 4.99, and *Dbh^LC-null^* mice (lower, 4.41; upper, 8.44). **, p=0.0067; ****, p<0.0001. GRAB_NE_ cohort: n=13 controls (6 F, 7 M), n=14 *Dbh^hypo^* mice (7 F, 7 M), n=11 *Dbh^LC-null^* (6 F, 5 M). GRAB_DA_ cohort: n=17 controls (8 F, 9 M), n=12 *Dbh^hypo^* mice (6 F, 6 M), n=12 *Dbh^LC-null^* (6 F, 6 M).

We next assessed NE and DA release in dCA1 during freezing in the recent (24 hours after shock) and remote (2 weeks after shock) tests which assess short- and long-term retrieval of the contextual fear memory. Both *Dbh^hypo^* (p=0.0384) and wild type controls (p=0.0003) exhibited a significant decrease in NE release during freezing in the recent test (Fig. 5A-B). As expected, we observed no change in NE release in *Dbh^LC-null^* mutants (Fig. 5A-B). DA release was also decreased during freezing in the recent test, however this decrease occurred in all genotypes including *Dbh^LC-null^* mice, and there was no significant difference found between the groups (Fig. 5 C-D). No significant sex-specific differences in NE or DA release were observed in the recent test. We observed no genotype-specific decrease in NE release during freezing in the remote test of long-term retrieval (Fig. 6 A-B), although male mice exhibited a significant decrease relative to females (p=0.0484, main effect of sex, 2-way ANOVA with Tukey’s test). Consistent with short-term retrieval, DA release was decreased in all mice during freezing in the remote test with no significant difference found between the groups (Fig. 6C-D). Altogether, these data identify unique patterns of NE and DA release dynamics in dCA1 during memory acquisition and retrieval that are differentially sensitive to partial or complete embryonic loss of NE.

**Figure 5.**
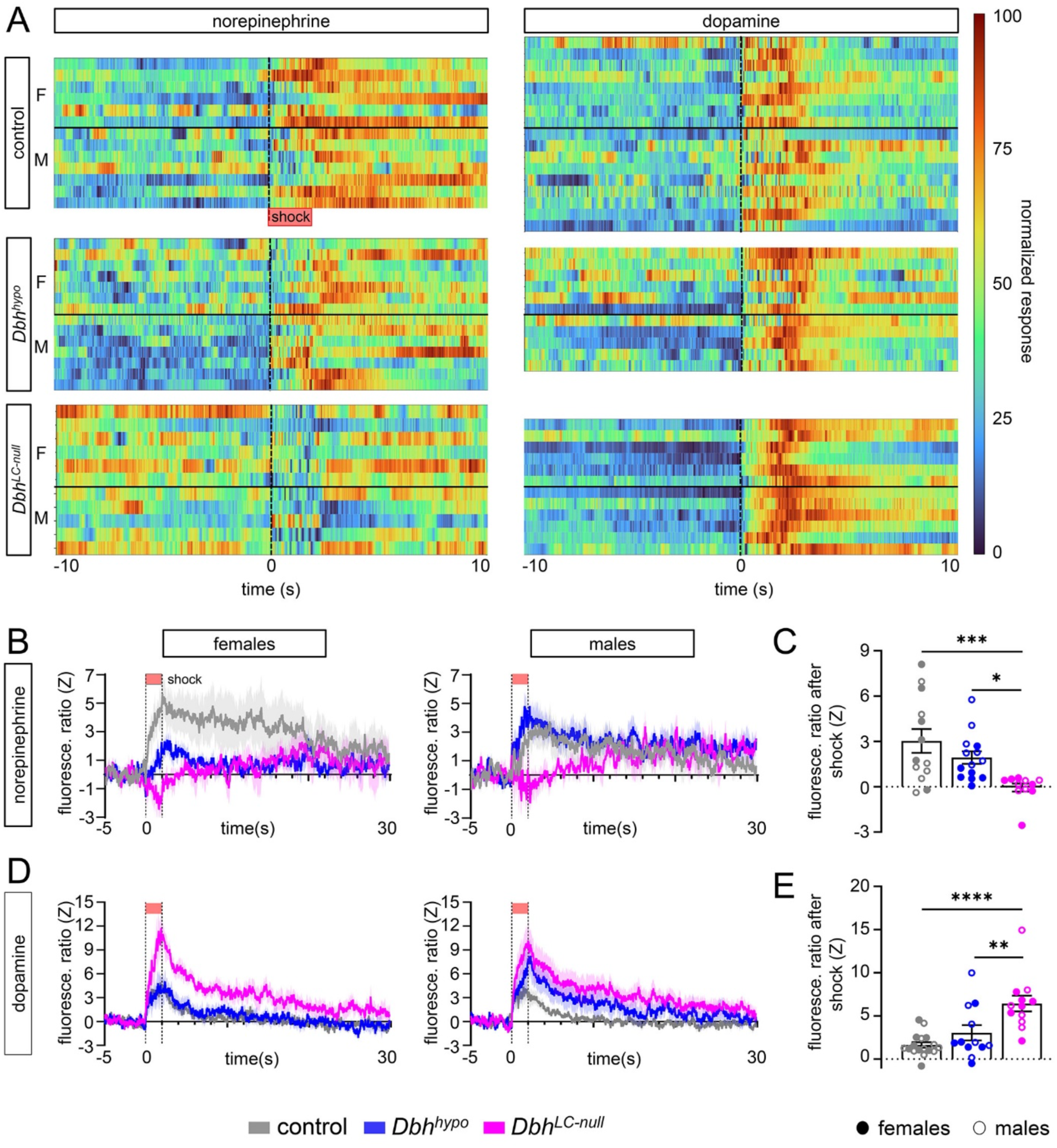
NE and DA dynamics in dCA1 during freezing in the recent context test. (**A**) Average GRAB_NE_/tdTomato fluorescence ratios aligned to freezing in female (left) and male (right) control, *Dbh^hypo^*, and *Dbh^LC-null^* mice. (**B**) Averages of NE response during the first six seconds of freezing episodes in controls (95% CI of mean: lower, -1.3; upper, -0.36), *Dbh^hypo^* (lower, -0.70; upper, -0.23), and *Dbh^LC-null^* mice (lower, -0.12; upper, 0.29; encompasses zero). **p=0.0003; *p=0.0384. (**C**) Average GRAB_DA_/tdTomato fluorescence ratios aligned to freezing. (**D**) Average DA responses during the first six seconds of freezing episodes in controls (95% CI of mean: lower, -0.61; upper, -0.18), *Dbh^hypo:^* (lower, -0.78; upper, -0.35), *Dbh^LC-null^* (lower, -0.77; upper, -0.14). Data are mean ± SEM. 2-way ANOVA with Tukey’s multiple comparisons test. GRAB_NE_ cohort: n= 13 controls, (6 F, 7 M), n= 14 *Dbh^hypo^* mice (7 F, 7 M), n= 11 *Dbh^LC-null^* (6 F, 5 M). GRAB_DA_ cohort: n= 18 *Dbh^wt/wt^* controls (9 F , 9 M), n= 12 *Dbh^hypo^* mice (6 F, 6 M), n= 12 *Dbh^LC-null^* (6 F, 6 M).

**Figure 6.**
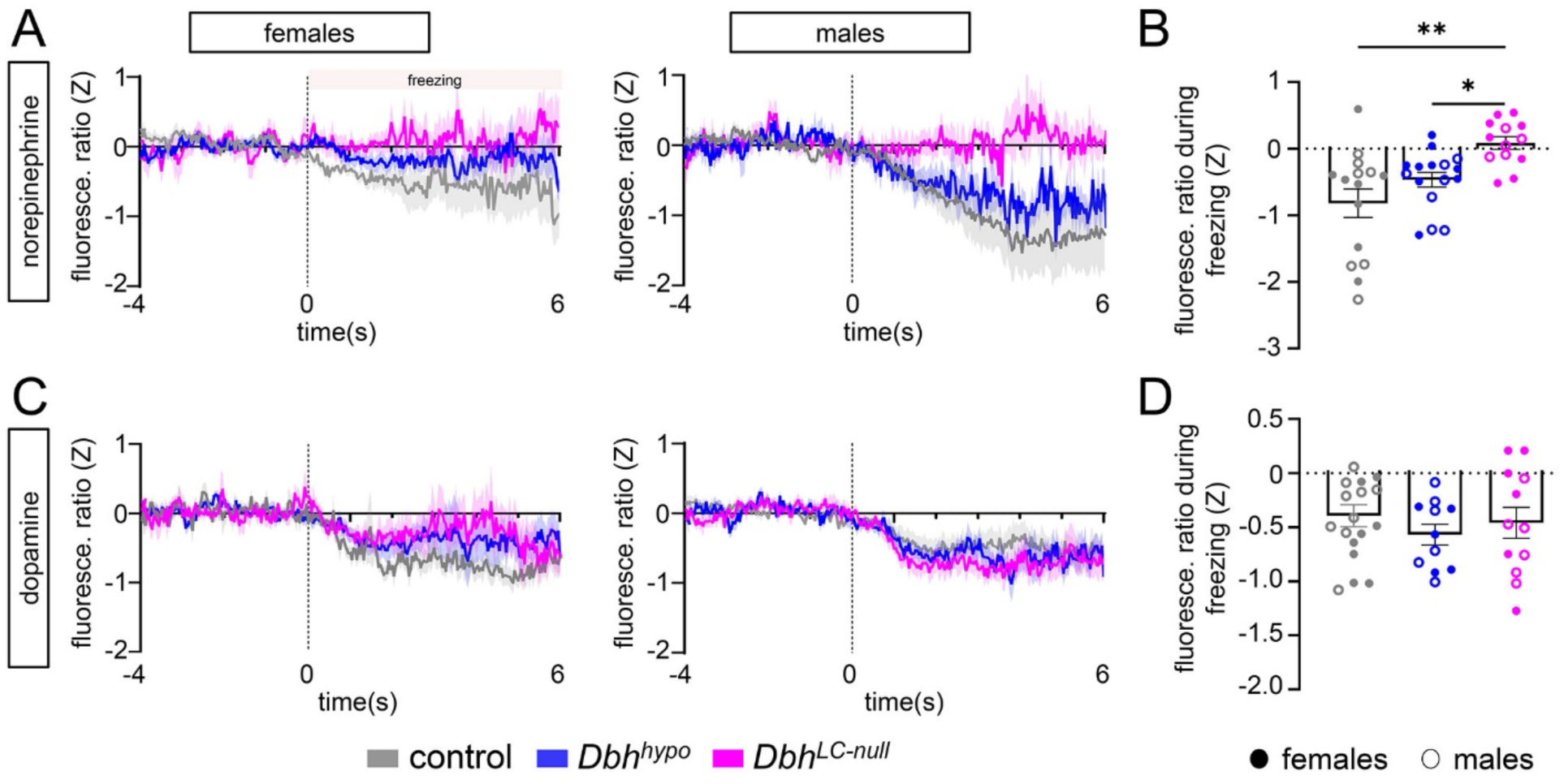
NE and DA dynamics in dCA1 during freezing in remote context test. (**A**) Average GRAB_NE_/tdTomato fluorescence ratios aligned to freezing in female (left) and male (right) control, *Dbh^hypo^*, and *Dbh^LC-null^* mice. (**B**) Average NE response during the first six seconds of freezing in control mice (95% CI of mean: lower, -0.64; upper, 0.057), *Dbh^hypo^* (lower, -0.43; upper, -0.06), and *Dbh^LC-null^* mice (lower, -0.25; upper, 0.13). (**C**) Average GRAB_DA_/tdTomato fluorescence ratios aligned to freezing. (**D**) Average DA responses during the first six seconds of freezing episodes in controls (95% CI of mean: lower, -0.78; upper, -0.29), *Dbh^hypo^* (lower,-0.53; upper,-0.063), and *Dbh^LC-null^* lower, -0.64; upper, -0.087). Data are mean ± SEM. 2-way ANOVA with Tukey’s multiple comparisons test. GRAB_NE_ cohort: n= 12 controls, (5 F, 7 M), n= 14 *Dbh^hypo^* (7 F, 7 M), n= 11 *Dbh^LC-null^* (6 F, 5 M). GRAB_DA_ cohort: n= 18 controls (9 F, 9 M), n= 12 *Dbh^hypo^* (6 F, 6 M), n= 12 *Dbh^LC-null^* (6 F, 6 M).

## DISCUSSION

Here we describe two new mouse models of disrupted *Dbh* expression beginning prenatally: a fortuitous hypomorphic allele, *Dbh^hypo^*, with reduced NE in central noradrenergic neurons and an LC conditional knockout, *Dbh^LC-null^*, with total loss of NE in the LC and reduced NE in other noradrenergic centers. We exploited these two models of disrupted NE synthesis to link catecholamine dynamics with specific outcomes in the context-dependent fear conditioning paradigm. Our data identify deficits in memory retrieval and unique patterns of catecholamine release dynamics in dCA1 during memory acquisition and retrieval that are differentially sensitive to partial or complete embryonic loss of NE.

We report that during acquisition of the contextual fear memory and the ten seconds immediately following the shock, NE and DA release was increased in dCA1 of both wild type and *Dbh^hypo^* mice. Although we found ∼61% reduction in NE and ∼113% increase in DA tissue content in the hippocampus of *Dbh^hypo^* mice, we observed no significant difference in either NE or DA release compared to wild type controls. DA release was only significantly increased in the total absence of NE release in dCA1 of *Dbh^LC-null^* mutants.

The absence of increased DA release in dCA1 of *Dbh^hypo^* mice, despite the significant increase in tissue content, highlights the tight regulation of release dynamics. The increase in DA tissue content was found in the hippocampus of both *Dbh* mutants, and was inversely proportional to deficits in NE content, suggesting that the increase is a direct result of decreased conversion of DA to NE in the LC. Several studies have shown that LC neurons are capable of releasing DA when artificially stimulated (34, 35), but other experiments suggest that midbrain dopaminergic neurons are the most likely natural source of DA in dorsal CA1 (14). Our data do not reveal the origin of the increased DA release observed in *Dbh^LC-null^* mice. Although it is plausibly due to an accumulation of DA in LC neurons, it could also be an indirect response of midbrain dopaminergic neurons to decreased LC-NE signaling.

The large release of NE in response to the foot-shock observed in wild type mice argues that the dorsal hippocampus is powerfully modulated by the aversive unconditional stimulus. While this observation is consistent with prior studies examining the LC response to noxious stimuli (36–41), it has important implications for theories of dorsal hippocampal involvement in contextual fear. The role of the dorsal hippocampus in forming an integrated multi-sensory representation of the context is typically emphasized, but not its role in representing the associative valence of the context, which is thought to be mediated by strengthening of synaptic connectivity between the contextual representation cells and downstream structures such as the amygdala (42). Modulation of the hippocampus by shock-induced NE release may serve to strengthen and/or modify the contextual representation, consistent with prior work that shock presentation causes re-mapping of place cells (43, 44). In addition, there is some evidence that the shock itself may become integrated into the contextual representation directly, which is thought to play an important role in fear extinction (45), a process that is strongly modulated by LC-NE signaling (40, 46).

In contrast to the large increase in NE release in dCA1 of wild type mice during memory acquisition, we observed decreased release during freezing in the test of short-term memory retrieval. Both *Dbh^LC-null^* and *Dbh^hypo^* mice had deficits in freezing; however, *Dbh^hypo^* mice exhibited decreased release of NE similar to wild type. In both *Dbh* mutants, DA release decreased similar to that of wild type mice. During freezing in the test of long-term memory retrieval we found that NE release was decreased in wild type and *Dbh^hypo^* males. We also observed a decrease in DA release in wild type and mutant males and females, with no significant genotype-specific differences, similar to freezing during short term retrieval. Thus, the deficits in freezing behavior do not appear to be directly correlated with loss of a dynamic reduction in NE or DA release.

The reduction in NE release during freezing could be a consequence of opioids released as part of the fear response (47, 48). Opioid signaling plays a critical role in the overall fear response – it produces conditional analgesia to suppress awareness of any immediate injuries so that the animal can deal with the threat at hand (49). Additionally, opioid signaling plays an important role in regulation of threat learning by dampening the effectiveness of an expected unconditional stimulus, effectively reducing the prediction error, which is evident in phenomena such as blocking (50, 51). Dampening of NE during freezing could play a role in reducing attention toward poor predictors of threat as has been demonstrated for opioid signaling in the nucleus accubmens (52). Recent work has delineated a descending, anti-nociceptive LC projection that contrasts with an ascending, aversive LC projection (53). The present finding of reduced NE in dCA1 release during freezing adds further complexity to this story and suggests a complex orchestration of functionality distinct LC-NE modules during fear expression.

The observed reduction in NE release is also interesting considering that increasing LC activity leads to expression of anxiety-related behaviors and is aversive (54, 55). A possible unifying explanation comes from findings that lower levels of tonic LC activity enhance signal detection and behavioral performance in attentionally demanding tasks (56). Higher tonic LC activity, in contrast, may facilitate evaluation of the sensory environment that is more appropriate in new and uncertain conditions (56). Thus, what we may be observing during expression of freezing is a shift from higher tonic activity during exploration of the environment to a lower level of tonic activity that enhances detection of threat (57).

Several notable sex differences were observed in this study that are worthy of discussion given the well-established sex differences in NE signaling (58). Male controls showed higher NE levels in the hippocampus, suggesting baseline sex differences in central NE production. The only overt sex difference observed in the behavioral measures was the greater distance moved by male mice during foot-shock. Interestingly, we also found a trend for a sex interaction in NE release in response to the foot shock whereby *Dbh^hypo^* males showed similar levels of NE release to the controls and *Dbh^hypo^* females more closely resembled *Dbh^LC-null^*. In addition, we observed a numerically greater decrease in NE release in male wild type and *Dbh^hypo^* mice compared to females during freezing in the remote test. These findings highlight the complex relationship between sex, genotype and NE signaling.

Comparisons of the *Dbh* mouse models used in the current study with a full-body *Dbh* knockout (*Dbh^ko^*; (22)) indicate that recall of fear memories is sensitive to levels of NE and the timing of the insult. *Dbh^ko^* mice have a deficit in recent retrieval of fear memories, but remote fear is intact (59), suggesting that NE is not required at remote time points when fear memories have already gone through a systems consolidation process. In the current study, deficits in retrieval of both recent and remote contextual fear memories found in *Dbh^LC-null^* mutants suggest a more critical role for LC-NE in remote retrieval. The difference in recall of remote fear memories between the three *Dbh* mutant models may be at least partially due to differences in NE levels during prenatal development. *Dbh^LC-null^* mice lack LC-NE throughout prenatal and postnatal development. Both *Dbh^hypo^* and *Dbh^KO^* mice have at least partial NE synthesis during embryonic development. In the case of the *Dbh^KO^* mice, low levels of NE synthesis result from prenatal L-DOPS treatment to avoid the embryonic lethal effect of total NE loss (22). Future work will be necessary to determine the requirement of NE during prenatal development in mediating the observed phenotypes.

## MATERIALS AND METHODS

### Animals

All procedures related to animal use were approved by the Animal Care and Use Committee (ACUC) of the National Institute of Environmental Health Sciences. Animal procedures were performed in accordance with the recommendations in the Guide for the Care and Use of Laboratory animals of the National Institutes of Health. All animals were group-housed and maintained on a 12-hour light/dark cycle at 72±2 °F with access to food and water *ad libitum*.

### Generation of a fluorescently tagged conditional knockout allele of *Dbh*

To generate the *Dbh^tm2.2Pjen^* (*Dbh^flx^*) allele, we first generated plasmid pGEM-loxrox-Hygro/Zeo (Fig. S2), consisting of a loxP site and rox-flanked hygromycin and zeocin resistance genes from pNeDaKO-Hyg (Addgene plasmid #16414; (60)). We also generated pGEM-2ATNSAG, consisting of a 2A peptide (26), tdTomato cDNA, SV40 poly(A) cassette, FRT-flanked neomycin/kanamycin resistance gene from pL451 (61), loxP, splice acceptor from pSA-βGeo (Addgene plasmid #21709; (62)), EGFP cDNA, and rabbit β-globin poly(A) (Fig. S2). Using homologous recombination in *Escherichia coli* strain DY380 (63), we inserted the loxrox-Hygro/Zeo and 2ATNSAG cassettes into BAC bMQ 351i22 (64) which contains the *Dbh* locus from a 129/SvEv mouse. Recombinant BACs were identified by ability to grow in the presence of zeocin and kanamycin and confirmed by polymerase chain reaction (PCR). Finally, we subcloned a 33.4 kb fragment from the modified BAC into pMCS-DTA (Addgene plasmid #22730; (65) to generate a targeting vector which extends from 4.9 kb upstream of the loxrox-Hygro/Zeo insertion to 5 kb downstream of the 2ATNSAG insertion.

The targeting vector was electroporated into G4 embryonic stem cells (B6129 F1 genetic background; (66)), and recombinant clones were selected by ability to grow in the presence of G418 and Hygromycin B. Homologous recombinants were confirmed by a combination of long-range PCR and Southern blotting before injection into B6(Cg)-*Tyr^c-2J^* blastocysts to produce chimeric mice. Male chimeras were bred to female C57BL/6J mice to establish the *Dbh^flx^* mouse line. We observed germline transmission of two ES clones and selected one for further analysis. Founding heterozygotes were initially crossed with B6;129-*Tg(CAG-dre)1Afst* (67) and B6.Cg-*Tg(ACTFlpe)9205Dym/J* mice (68) to excise the rox- and FRT-flanked antibiotic resistance cassettes, and the line was subsequently maintained by back-crossing to C57BL/6J.

### Experimental crosses

To confirm that the loxP sites in the *Dbh^flx^* allele are recombined by Cre recombinase, resulting in deletion of exons 2-12, we crossed *Dbh^flx^* heterozygotes with C57BL/6J.Cg-*Tmem163^Tg(ACTB-cre)2Mrt^* mice (27). To generate *Dbh^LC-null^* mice, we first intercrossed *Dbh^flx^* and C57BL/6J.Cg-*En1^tm2(cre)Wrst^* (*En1^Cre^*) heterozygotes (28) and then backcrossed double heterozygous offspring to *Dbh^flx^* heterozygotes or homozygotes.

### Genotyping and genomic PCR

Mice were routinely genotyped using automated real-time PCR (Transnetyx, Cordova, TN). Screening included an assay to detect rare germline recombination of *Dbh^flx^* associated with *En1^Cre^* expression. To distinguish *Dbh^wt^*, *Dbh^flx^*, and fully recombined *Dbh^null^* alleles by agarose gel electrophoresis (Fig. 1A), we used primers (i) 5’-GACAGAGCAGGGCTGGCTGTACAG, (ii) 5’-TAGCAACATTTCCTCTAGGGAC, (iii) 5’-TAATCAGCCATACCACATTTGTAGAG, and (iv) 5’- ACAGGATAAGTATGACATCATCAAGG.

### Tissue/plasma collection

For immunohistochemistry, mice were anesthetized with sodium pentobarbital and transcardially perfused with 4% PFA. Brains were dissected and post-fixed in 4% PFA at 4 °C overnight. Following a rinse in 1X PBS, brains were equilibrated in 30% sucrose/PBS for 48 hours at 4 °C and embedded in Tissue Freezing Medium (General Data Company, Cincinnati, OH). Brains were cryosectioned (40-µm free-floating sections) and sections stored at -80 °C in 30% sucrose/30% ethylene glycol in 1X PBS.

For catecholamine analysis, 100-200 µl of whole blood was collected in EDTA-treated tubes using a submandibular blood collection procedure. Collected blood was centrifuged for 10 minutes at 2,000 x g, and supernatant (100-200 µl) was flash-frozen in liquid nitrogen and stored at -80 °C. For detection of brain catecholamines, mice were euthanized by CO_2_ inhalation. Hippocampal tissue was rapidly dissected, flash-frozen in liquid nitrogen and stored at -80 °C. Mass spectrometry detection of catecholamines and metabolites was performed by the Vanderbilt University Neurochemistry Core Laboratory.

To obtain separate RNA samples from noradrenergic populations in pons and medulla, mice were euthanized with an overdose of sodium pentobarbital, and the hindbrain was isolated. The hindbrain was cut sagittally at the midline and one half was bisected in the dorsoventral plane from the rostral edge of the obex to the caudal edge of the pontine gray. The cerebellum was removed and the pons and medulla were flash-frozen in liquid nitrogen and stored at -80 °C prior to isolation of RNA.

### Immunohistochemistry

For immunofluorescent labeling of noradrenergic neurons, the following primary antibodies were used: rabbit anti-dsRed (1:1,000, Cat.# 632496, Clontech Laboratories, Mountain View, CA), chicken anti-GFP (1:10,000, Cat.# ab13970, Abcam, Cambridge, MA), mouse anti-tyrosine hydroxylase (TH) (1:500, clone 185, Cat.# GTX10372, GeneTex, Irvine, CA), and rabbit anti-DBH (1:500, Cat.# SAB2701977, Sigma Aldrich, St. Louis, MO). Secondary Antibodies (all used at 1:1000 dilution) were Alexa Fluor 568 goat anti-rabbit (Cat.# A11036, Thermofisher Scientific, Waltham, MA), Alexa Fluor 488 goat anti-chicken (Cat.# A11039, Thermofisher Scientific), Alexa Fluor 633 goat anti-mouse (Cat.# A21052, Thermofisher Scientific), Alexa Fluor 568 goat anti-mouse (Cat.# A11031, Thermofisher Scientific), and Alexa Fluor 488 goat anti-rabbit (Cat.# A11034, Thermofisher Scientific. Immunolabeled brain sections were stained with 4^′^, 6- diamidino-2- phenylindole (DAPI) or Neurotrace 435/455 Blue Fluorescent Nissl stain (Thermofisher Scientific), mounted on Superfrost Plus slides, and coverslipped with VectaShield Hard Set mounting medium (Vector Laboratories, Burlingame, CA).

### Digital image acquisition and processing

Tile-scan images were collected on Zeiss 780 or 880 inverted confocal microscopes (Carl Zeiss Microscopy, Thornwood, NY) using a 20X or 40X objective. Z-stacks were converted to maximum intensity projections with Zen 2012 Black Software (Carl Zeiss). Images were modified only by adjusting brightness and contrast across the entire image in Photoshop (Adobe Systems, San Jose, CA) or ImageJ software (US National Institute of Health). Anatomical location was confirmed by reference to a mouse brain atlas (69).

### Droplet digital PCR (ddPCR)

Frozen tissue was placed in 500 µl Trizol Reagent (ThermoFisher Scientific) and homogenized using a battery-powered Pellet Mixer and disposable pestle (VWR International, Radnor, PA). After chloroform extraction, the aqueous layer was mixed with an equal volume of 75% ethanol and transferred to an RNeasy spin-column (Qiagen, Germantown, MD). RNA cleanup and elution was performed according to manufacturer’s instructions for the RNeasy Mini kit. Concentration and integrity of eluted RNA was evaluated using a Fragment Analyzer (Agilent Technologies, Santa Clara, CA), and RNA was stored at -80 °C.

For ddPCR, 250 ng total RNA was reverse transcribed using the iScript cDNA synthesis kit (BioRad, Hercules, CA). ddPCR probes (BioRAd) were *Dbh* (dMmuCPE5117134, FAM fluorophore), *Th* (dMmuCPE5121062, FAM), and *Tbp* (dMmuCPE5124759, HEX). PCR reactions were assembled using the ddPCR Supermix for Probes (No dUTP) kit (BioRad). Each ddPCR reaction contained 1 µl cDNA and either *Dbh* and *Tbp* or *Th* and *Tbp* probes (250 nM each) in a final volume of 25 µl. Twenty microliters of each reaction mix was converted to droplets using the QX200 Droplet Generator (BioRad), transferred to a new 96-well plate, sealed with PX1 PCR plate sealer (BioRad) and cycled in a T100 Thermal Cycler (BioRad) under the following protocol: 40 cycles of 95 °C for 30 seconds and 68 °C for 1 minute, followed by a single cycle of 98 °C for 10 minutes. The plate was then read in the FAM and HEX channels using the QX200 Droplet Reader, and the concentration of each target RNA (template copies per microliter of total RNA) was estimated using Quanta-Soft ddPCR software (BioRad).

For each RNA sample, three replicate reactions were performed, and the results of the replicates were averaged. *Dbh* and *Th* expression data were normalized by *Tbp* expression. A normalization factor was calculated for each sample by dividing *Tbp* concentration by the average *Tbp* concentration of all samples. Normalized *Dbh* and *Th* concentration was then calculated by dividing the raw concentration by the normalization factor for that sample.

### Surgery

For viral injection and fiber optic implantation, mice were anesthetized with 4% isofluorane and placed in a Model 900 stereotaxic apparatus (Kopf Instruments, Los Angeles, CA) equipped with a digital micromanipulator (Harvard Apparatus, Holliston, MA). Bupivicaine (270 µg in 0.1ml) was injected locally beneath the scalp prior to incision, and anesthesia was maintained with 0.5-2% isoflurane during the surgery. Hippocampal area CA1 was injected with a GRAB_NE_ NE sensor (AAV9-hSyn-GRAB_NE1m; Addgene #123308; (32)) or GRAB_DA_ DA sensor (AAV9-hSyn-GRAB-DA2m; Addgene # 140553; (33)) using the following stereotaxic coordinates: −2.00 posterior, −1.25 lateral, −1.25 ventral (mm from Bregma). Each sensor was co-injected with tdTomato (AAV5-hSyn-tdTomato) at 1:5 ratio. AAVs (500 nL) were delivered at 100-nL/minute using a 30-gauge Neuros syringe (Hamilton Company, Reno, NV) and UMP3 UltraMicroPump (World Precision Instruments, Sarasota, FL), and needles were left in place for 5 minutes after infusion to minimize backflow of the virus upon withdrawal. All mice received the analgesic Buprenorphine SR (1 mg kg^-1^ s.c.) after surgery.

Three weeks after the viral injections, mice were unilaterally implanted with optical probes above dCA1 (−2 posterior, −1.40 lateral, −1.35 ventral, mm from bregma). The optical probes, made from 0.39 NA, 200-µm core multimode optical fiber (FT200EMT; Thorlabs, Newton, NJ), were adhered with epoxy (PFP 353ND; Precision Fiber Products, Chula Vista, CA) within a 1.25-mm OD ceramic ferrule (MM-CON2007–2300 or MM-CON2010-1270-2-WHT; Precision Fiber Products). Final placement of the optical fiber probe was guided by live fluorescence signal. To accomplish this, excitation light from a 470 nm, 760 mW mounted LED (M470L4, Thorlabs) was launched into a DFM1 fluorescence cube (Thorlabs) containing an ET470/40x excitation filter and T495lpxt beam splitter (Chroma Technology Corp., Rockingham, VT), reflected into an achromatic fiber port (PAF2-A4A, Thorlabs), and collected by a 5-meter-long relay multimode patch cable (M72L05, 200-µm core with 0.39 NA, Thorlabs). The patch cable was connected to the implanted optical fiber via a second 1-meter-long multimode patch cable (M83L01: 200-µm core with 0.39 NA or M61L01: 105-µm core with 0.22 NA, Thorlabs) which was linked to the ferrule containing the optical fiber by a ceramic sleeve (SMCS125S, Precision Fiber Products). Excitation light from the implantable fiber was adjusted to ∼70-µW. Emitted light was passed through an emission filter (ET500lp, Chroma Technology Corp.), collected into an Ocean FX spectrometer (Ocean Insight, Orlando, FL), and visualized using Ocean View version 1.6.7 (Ocean Insight). When final placement was confirmed by detection of large tdTomato and GRAB_NE_ or GRAB_DA_ fluorescent peaks, the optical probe was anchored to the skull using C&B Metabond (Parkell, Edgewood, NY) and dental acrylic (Lang Dental Manufacturing, Wheeling, IL). Mice were given at least 1-week for surgical recovery before habituation and behavioral testing.

### Fear Conditioning

For each day of the procedure, animals were brought by cart to a holding area where they were left untouched in their home cages for at least 30 minutes. Two 15-minute habituation sessions over two days accustomed the mouse to being handled, anesthetized, and connected to the patch cable. Each mouse was briefly anesthetized with isoflurane, connected to a patch cable, and allowed rest in a clean cage for 15 minutes in a red-light room with an overhead light producing 35 lux. On test days, the mouse was then transferred to a fear conditioning chamber with grid floor (Med Associates, Fairfax, VT) for the experimental procedure. *Pre-Exposure:* the mouse was allowed to explore the conditioning chamber for 10 minutes. *Immediate Shock Training (24 hours later):* Ten seconds after the mouse was placed in the conditioning chamber, a 2-second, 0.75 mA shock was presented. The mouse was removed from the chamber 30 seconds after shock for a total of 42 seconds in the conditioning chamber*. Recent Context Test (24 hours later) and Remote Context Test (2 weeks later):* The mouse was returned to the conditioning chamber for 10 minutes, during which freezing behavior was quantified. Between mice, the chamber was cleaned with 70% ethanol, and the floor trays were rinsed with water then sprayed with Windex.

### Behavioral Tracking

Video was recorded using Video Freeze software (Med Associates), and distance moved (pre-exposure, immediate shock training) and freezing (pre-exposure, recent and remote context tests) were tracked. Pixel noise generated by the patch cable prevented the use of the Video Freeze automated freeze detection, so we validated an alternative method using Ethovision XT software vs. 16 (Noldus Information Technology, Leesburg, VA) in which movement tracking was limited to a smaller arena within the video which contained the mouse but not the patch cable. Parameters for detecting freezing in Ethovision were validated against three human observers (Fig. S3).

### *in vivo* Photometry Recordings

GRAB_NE_, GRAB_DA_, and tdTomato fluorescence intensities were measured in dCA1 neurons using a custom-built fiber photometry system, as previously described (31). Excitation light from a 488 nm, 20 mW continuous wave laser (OBIS 488LS-20; Coherent, Santa Clara, CA) was launched into a DFM1 fluorescence cube, reflected by a dichroic mirror (ZT488/561rpc-UF1; Chroma Technology Corp.), and focused by an achromatic fiber port (PAFA-X-4-A; Thorlabs) onto the core of a multimode patch cable (M83L01; Thorlabs). The distal end of the patch cable was connected to the implanted optical probe by a quick-release interconnect (ADAL3; ThorLabs), and the power of the excitation light was adjusted to 66 μW. Emitted fluorescence was collected by the optical fiber probe and patch cable, passed through the dichroic mirror, and filtered through an emission filter (ZET 488/561m; Chroma Technology Corp.) before being collected by a fiber port (PAF2S-11A; Thorlabs) and launched into a QE Pro-FL spectrometer (Ocean Insight) through a multi-mode patch cable (M200L02-A, 0.22 NA, AR-coated, 200/240 μm core/cladding; Thorlabs). Time-lapsed fluorescence emission spectra were visualized using OceanView version 1.5 (Ocean Insight). The spectrometer and fear conditioning chamber were triggered using a Doric TTL pulse generator (OTPG_4; Doric Lenses, Quebec, Canada) controlled by Neuroscience Studio software (Doric Lenses).

### Photometry Data Analysis

To quantify GRAB_NE_ or GRAB_DA_ fluorescence, and to separate spectral overlap between the sensor and tdTomato, all raw emission spectra data were passed through a spectral linear unmixing algorithm written in R, as described previously (31). To control for movement artifacts in the fluorescence signal (e.g. photon loss caused by tissue movement or bending of the fiber during mouse movement), the unmixed GRAB_NE_ or GRAB_DA_ coefficients were normalized to unmixed tdTomato coefficients to generate GRAB_NE_/tdTomato or GRAB_DA_/tdTomato fluorescence ratios.

Behavioral-event-triggered averages were used to analyze the photometry data across groups using the custom coded fiber photometry analysis script (FiPhA), coded in R ((70); Fig. S4). Using FiPhA to analyze experimenter-defined events, such as the foot shock recordings, the raw fluorescent ratio values were aligned to the start of the shock and z-scores (relative to the 4 second baseline) of the raw photometry ratios were calculated. Aligned photometry traces were averaged across genotype and male and female differences were plotted. To compare the signals across groups, the z-scored fluorescent ratio values were averaged for each group for ten seconds after the foot shock and plotted (Fig. S4A).

For the whole recording session, behavioral freezing episodes in the recent (24-hour) and remote (2-week) tests were also aligned with the raw GRAB_NE_/tdTomato or GRAB_DA_/tdTomato fluorescent ratios. To allow for adequate baseline recordings, any freezing episodes that occurred within five seconds of the recording session start were dropped, and any episodes that occurred within five seconds of each other were aggregated. Freezing episodes shorter than one second were excluded. All freezing episodes within the recording sessions were aligned and the raw photometry ratios were z-scored, with a four second baseline value, and averaged event traces were plotted across genotype and sex. To compare the NE and DA recordings across genotypes, z-scored values were averaged for six seconds during a freezing event and the values were plotted (Fig S4B-E).

### Statistical Analysis

For comparison of experimental groups, we used GraphPad Prism 9 software to perform a 2-way ANOVA with genotype and sex as factors followed by Tukey’s multiple comparison test when justified (Dataset S1). Where sex differences were observed within a group and statistical comparisons are justified by a main effect of sex or genotype-by-sex interaction, this is indicated with ### in the figures and/or described in the main text. No statistically significant differences were observed between *En1^Cre/wt^*; *Dbh^wt/wt^* and *En1^wt/^*^wt^; *Dbh^wt/wt^* mice in behavioral assays and measurements of NE and DA levels, so they were graphed together as controls (Fig. S5).

## ACKNOWLEDGEMENTS

We would like to thank Sandra McBride and Matthew Bridge (DLH corporation) for the custom R code used to analyze photometry data; Wesley Gladwell and Kevin Gerrish (NIEHS Molecular Genomics Core Facility) for performing ddPCR; and the Vanderbilt Neurochemistry Core for catecholamine mass spectometry. Valuable support was also provided by the NIEHS Comparative Medicine Branch, and the Neurobehavioral, Fluorescence Microscopy and Imaging, and Viral Vector Cores. We would also like to thank Dr. Guohong Cui for valuable discussion of the photometry data. This research was supported by the Intramural Research Program at the National Institute of Environmental Health Sciences (ZIA ES102805 to P.J., and ZIC ES103330) and the Extramural Research Program at the National Institute of Diabetes and Digestive and Kidney Diseases (R00 DK119586 to N.R.S.).

## Author contribitions

N.W.P., J.D.C. and P.J. designed research; L.R.W., N.W.P., I.Y.E., D. P., C.L.S, K.G.S., S.A.F., A.L.D., V.W.K., and P.J. performed research; L.R.W., N.W.P., I.Y.E., N.R.S., J.D.C., and P.J. analysed data; and L.R.W., N.W.P., J.D.C., and P.J. wrote the paper.

## REFERENCES

1. T. Bast, Toward an integrative perspective on hippocampal function: from the rapid encoding of experience to adaptive behavior. Rev Neurosci 18, 253–281 (2007).

2. J. W. Rudy, R. C. O’Reilly, Contextual fear conditioning, conjunctive representations, pattern completion, and the hippocampus. Behav Neurosci 113, 867–880 (1999).

3. D. M. Smith, D. A. Bulkin, The form and function of hippocampal context representations. Neurosci Biobehav Rev 40, 52–61 (2014).

4. J. D. Cushman, M. S. Fanselow, “Context Fear Learning” in Encyclopedia of the Sciences of Learning, N. M. Seel, Ed. (Springer US, Boston, MA, 2012), 10.1007/978-1-4419-1428-6_832, pp. 797–799.

5. F. B. Krasne, J. D. Cushman, M. S. Fanselow, A Bayesian context fear learning algorithm/automaton. Front Behav Neurosci 9, 112 (2015).

6. M. J. Sanders, B. J. Wiltgen, M. S. Fanselow, The place of the hippocampus in fear conditioning. Eur J Pharmacol 463, 217–223 (2003).

7. J. J. Kim, M. S. Fanselow, Modality-specific retrograde amnesia of fear. Science 256, 675–677 (1992).

8. R. G. Phillips, J. E. LeDoux, Differential contribution of amygdala and hippocampus to cued and contextual fear conditioning. Behav Neurosci 106, 274–285 (1992).

9. A. Chowdhury et al., A locus coeruleus-dorsal CA1 dopaminergic circuit modulates memory linking. Neuron 110, 3374–3388.e3378 (2022).

10. T. F. Giustino, S. Maren, Noradrenergic Modulation of Fear Conditioning and Extinction. Front Behav Neurosci 12, 43 (2018).

11. C. F. Murchison et al., A distinct role for norepinephrine in memory retrieval. Cell 117, 131–143 (2004).

12. D. O. Seo et al., A locus coeruleus to dentate gyrus noradrenergic circuit modulates aversive contextual processing. Neuron 109, 2116–2130.e2116 (2021).

13. W. Shen et al., Astroglial adrenoreceptors modulate synaptic transmission and contextual fear memory formation in dentate gyrus. Neurochem Int 143, 104942 (2021).

14. T. Tsetsenis et al., Midbrain dopaminergic innervation of the hippocampus is sufficient to modulate formation of aversive memories. Proc Natl Acad Sci U S A 118 (2021).

15. T. Tsetsenis, J. I. Broussard, J. A. Dani, Dopaminergic regulation of hippocampal plasticity, learning, and memory. Front Behav Neurosci 16, 1092420 (2022).

16. A. Wagatsuma et al., Locus coeruleus input to hippocampal CA3 drives single-trial learning of a novel context. Proc Natl Acad Sci U S A 115, E310–e316 (2018).

17. N. W. Plummer et al., Expanding the power of recombinase-based labeling to uncover cellular diversity. Development 142, 4385–4393 (2015).

18. S. D. Robertson, N. W. Plummer, J. de Marchena, P. Jensen, Developmental origins of central norepinephrine neuron diversity. Nat Neurosci 16, 1016–1023 (2013).

19. H. S. Lee, J. H. Han, Activity Patterns of Individual Neurons and Ensembles Correlated with Retrieval of a Contextual Memory in the Dorsal CA1 of Mouse Hippocampus. J Neurosci 43, 113–124 (2023).

20. K. Z. Tanaka et al., Cortical representations are reinstated by the hippocampus during memory retrieval. Neuron 84, 347–354 (2014).

21. T. Tsetsenis, J. K. Badyna, R. Li, J. A. Dani, Activation of a Locus Coeruleus to Dorsal Hippocampus Noradrenergic Circuit Facilitates Associative Learning. Front Cell Neurosci 16, 887679 (2022).

22. S. A. Thomas, A. M. Matsumoto, R. D. Palmiter, Noradrenaline is essential for mouse fetal development. Nature 374, 643–646 (1995).

23. J. W. Rudy, N. C. Huff, P. Matus-Amat, Understanding contextual fear conditioning: insights from a two-process model. Neurosci Biobehav Rev 28, 675–685 (2004).

24. D. L. Stote, M. S. Fanselow, NMDA receptor modulation of incidental learning in Pavlovian context conditioning. Behav Neurosci 118, 253–257 (2004).

25. S. A. Thomas, B. T. Marck, R. D. Palmiter, A. M. Matsumoto, Restoration of norepinephrine and reversal of phenotypes in mice lacking dopamine beta-hydroxylase. J Neurochem 70, 2468–2476 (1998).

26. G. Trichas, J. Begbie, S. Srinivas, Use of the viral 2A peptide for bicistronic expression in transgenic mice. BMC Biol 6, 40 (2008).

27. M. Lewandoski, E. N. Meyers, G. R. Martin, Analysis of Fgf8 gene function in vertebrate development. Cold Spring Harb Symp Quant Biol 62, 159–168 (1997).

28. R. A. Kimmel et al., Two lineage boundaries coordinate vertebrate apical ectodermal ridge formation. Genes Dev 14, 1377–1389 (2000).

29. C. R. Benedict, M. Fillenz, C. Stanford, Changes in plasma noradrenaline concentration as a measure of release rate. Br J Pharmacol 64, 305–309 (1978).

30. F. Cunningham, et al., Ensembl 2022. Nucleic Acids Res 50, D988–d995 (2022).

31. C. Meng et al., Spectrally Resolved Fiber Photometry for Multi-component Analysis of Brain Circuits. Neuron 98, 707–717.e704 (2018).

32. J. Feng et al., A Genetically Encoded Fluorescent Sensor for Rapid and Specific In Vivo Detection of Norepinephrine. Neuron 102, 745–761.e748 (2019).

33. F. Sun et al., Next-generation GRAB sensors for monitoring dopaminergic activity in vivo. Nat Methods 17, 1156–1166 (2020).

34. K. A. Kempadoo, E. V. Mosharov, S. J. Choi, D. Sulzer, E. R. Kandel, Dopamine release from the locus coeruleus to the dorsal hippocampus promotes spatial learning and memory. Proc Natl Acad Sci U S A 113, 14835–14840 (2016).

35. T. Takeuchi et al., Locus coeruleus and dopaminergic consolidation of everyday memory. Nature 537, 357–362 (2016).

36. J. M. Cedarbaum, G. K. Aghajanian, Activation of locus coeruleus neurons by peripheral stimuli: modulation by a collateral inhibitory mechanism. Life Sci 23, 1383–1392 (1978).

37. D. M. Devilbiss, C. W. Berridge, Low-dose methylphenidate actions on tonic and phasic locus coeruleus discharge. J Pharmacol Exp Ther 319, 1327–1335 (2006).

38. M. J. Mana, A. A. Grace, Chronic cold stress alters the basal and evoked electrophysiological activity of rat locus coeruleus neurons. Neuroscience 81, 1055–1064 (1997).

39. R. M. Neves, S. van Keulen, M. Yang, N. K. Logothetis, O. Eschenko, Locus coeruleus phasic discharge is essential for stimulus-induced gamma oscillations in the prefrontal cortex. J Neurophysiol 119, 904–920 (2018).

40. A. Uematsu et al., Modular organization of the brainstem noradrenaline system coordinates opposing learning states. Nat Neurosci 20, 1602–1611 (2017).

41. R. J. Valentino, S. L. Foote, Corticotropin-releasing hormone increases tonic but not sensory-evoked activity of noradrenergic locus coeruleus neurons in unanesthetized rats. J Neurosci 8, 1016–1025 (1988).

42. R. L. Redondo et al., Bidirectional switch of the valence associated with a hippocampal contextual memory engram. Nature 513, 426–430 (2014).

43. M. A. Moita, S. Rosis, Y. Zhou, J. E. LeDoux, H. T. Blair, Putting fear in its place: remapping of hippocampal place cells during fear conditioning. J Neurosci 24, 7015–7023 (2004).

44. J. Ormond, S. A. Serka, J. P. Johansen, Enhanced reactivation of remapping place cells during aversive learning. J Neurosci 10.1523/jneurosci.1450-22.2022 (2023).

45. A. F. Lacagnina et al., Distinct hippocampal engrams control extinction and relapse of fear memory. Nat Neurosci 22, 753–761 (2019).

46. T. F. Giustino, K. R. Ramanathan, M. S. Totty, O. W. Miles, S. Maren, Locus Coeruleus Norepinephrine Drives Stress-Induced Increases in Basolateral Amygdala Firing and Impairs Extinction Learning. J Neurosci 40, 907–916 (2020).

47. M. J. Christie, Mechanisms of opioid actions on neurons of the locus coeruleus. Prog Brain Res 88, 197–205 (1991).

48. N. M. Enman, B. A. S. Reyes, Y. Shi, R. J. Valentino, E. J. Van Bockstaele, Sex differences in morphine-induced trafficking of mu-opioid and corticotropin-releasing factor receptors in locus coeruleus neurons. Brain Res 1706, 75–85 (2019).

49. M. S. Fanselow, Conditioned fear-induced opiate analgesia: a competing motivational state theory of stress analgesia. Ann N Y Acad Sci 467, 40–54 (1986).

50. J. P. Johansen, J. W. Tarpley, J. E. LeDoux, H. T. Blair, Neural substrates for expectation-modulated fear learning in the amygdala and periaqueductal gray. Nat Neurosci 13, 979–986 (2010).

51. G. P. McNally, M. Pigg, G. Weidemann, Blocking, unblocking, and overexpectation of fear: a role for opioid receptors in the regulation of Pavlovian association formation. Behav Neurosci 118, 111–120 (2004).

52. M. D. Iordanova, G. P. McNally, R. F. Westbrook, Opioid receptors in the nucleus accumbens regulate attentional learning in the blocking paradigm. J Neurosci 26, 4036–4045 (2006).

53. S. Hirschberg, Y. Li, A. Randall, E. J. Kremer, A. E. Pickering, Functional dichotomy in spinal- vs prefrontal-projecting locus coeruleus modules splits descending noradrenergic analgesia from ascending aversion and anxiety in rats. Elife 6 (2017).

54. J. G. McCall et al., CRH Engagement of the Locus Coeruleus Noradrenergic System Mediates Stress-Induced Anxiety. Neuron 87, 605–620 (2015).

55. N. R. Sciolino et al., Recombinase-Dependent Mouse Lines for Chemogenetic Activation of Genetically Defined Cell Types. Cell Rep 15, 2563–2573 (2016).

56. M. Usher, J. D. Cohen, D. Servan-Schreiber, J. Rajkowski, G. Aston-Jones, The role of locus coeruleus in the regulation of cognitive performance. Science 283, 549–554 (1999).

57. K. Roelofs, P. Dayan, Freezing revisited: coordinated autonomic and central optimization of threat coping. Nat Rev Neurosci 23, 568–580 (2022).

58. D. A. Bangasser, K. R. Wiersielis, S. Khantsis, Sex differences in the locus coeruleus-norepinephrine system and its regulation by stress. Brain Res 1641, 177–188 (2016).

59. C. F. Murchison, K. Schutsky, S. H. Jin, S. A. Thomas, Norepinephrine and ß₁-adrenergic signaling facilitate activation of hippocampal CA1 pyramidal neurons during contextual memory retrieval. Neuroscience 181, 109–116 (2011).

60. M. Kohli, C. Rago, C. Lengauer, K. W. Kinzler, B. Vogelstein, Facile methods for generating human somatic cell gene knockouts using recombinant adeno-associated viruses. Nucleic Acids Res 32, e3 (2004).

61. P. Liu, N. A. Jenkins, N. G. Copeland, A highly efficient recombineering-based method for generating conditional knockout mutations. Genome Res 13, 476–484 (2003).

62. G. Friedrich, P. Soriano, Promoter traps in embryonic stem cells: a genetic screen to identify and mutate developmental genes in mice. Genes Dev 5, 1513–1523 (1991).

63. E. C. Lee et al., A highly efficient Escherichia coli-based chromosome engineering system adapted for recombinogenic targeting and subcloning of BAC DNA. Genomics 73, 56–65 (2001).

64. D. J. Adams et al., A genome-wide, end-sequenced 129Sv BAC library resource for targeting vector construction. Genomics 86, 753–758 (2005).

65. S. X. Chen et al., Quantification of factors influencing fluorescent protein expression using RMCE to generate an allelic series in the ROSA26 locus in mice. Dis Model Mech 4, 537–547 (2011).

66. S. H. George et al., Developmental and adult phenotyping directly from mutant embryonic stem cells. Proc Natl Acad Sci U S A 104, 4455–4460 (2007).

67. K. Anastassiadis et al., Dre recombinase, like Cre, is a highly efficient site-specific recombinase in E. coli, mammalian cells and mice. Dis Model Mech 2, 508–515 (2009).

68. C. I. Rodríguez et al., High-efficiency deleter mice show that FLPe is an alternative to Cre-loxP. Nat Genet 25, 139–140 (2000).

69. G. Paxinos, K. B. J. Franklin, The Mouse Brain in Stereotaxic Coordinates (Academic press, New York, ed. 4th, 2013).

70. N. R. Sciolino et al., Natural locus coeruleus dynamics during feeding. Sci Adv 8, eabn9134 (2022).

